# Receptive-field sizes during remapping and uniform transsaccadic updating across the visual space

**DOI:** 10.64898/2026.05.25.727689

**Authors:** Yiming Wang, Mingsha Zhang, Ning Qian

## Abstract

Forward receptive-field (RF) remapping, a mechanism for transsaccadic updating of retinal positions and perceptual stability, transiently changes cells’ eccentricities and thus could also change their RF sizes, yet few studies examined RF sizes during remapping. A related issue is how the mechanism ensures the desired uniform updating across the visual space – a subtraction of the saccade vector from stimuli’s retinal positions wherever they appear – given highly nonuniform RF sizes and cortical magnification over eccentricities. We analyzed our recent circuit model for remapping/updating after incorporating eccentricity-dependent RF sizes and found that when the corollary-discharge-gated connections achieve uniform updating in the visual space, the model predicts no change to cells’ RF sizes despite their receiving inputs from other cells with different RF sizes. In contrast, if the updating were uniform in the cortical space but not visual space, cells’ RF sizes would change during remapping. We analyzed the data from the lateral intraparietal area and frontal eye fields and found that remapping magnitudes are similar for cells of different eccentricities. We then confirmed the prediction that RF sizes did not change significantly during remapping. These results reveal a circuit mechanism for uniform updating and perceptual stability across the entire visual field.

## Introduction

Forward RF remapping, also known as predictive remapping, is the observation that around the time of a saccade, some cells’ RFs (perisaccadic RFs or pRFs) shift in the saccade direction, driven by the corollary discharge (CD) of the saccade command ^1–6^. Early studies showed that the lateral intraparietal area (LIP) and frontal eye fields (FEF) have cells that respond to stimuli in their future (postsaccadic) RF (fRF) locations, accompanied by reduced responses to stimuli in their current (presaccadic) RF (cRF) locations, even before the saccade onset. (A cell’s cRF and fRF are its classic, retinotopic RF on the screen well before and well after the saccade, respectively; they superimpose on the retina.) Recent work measured the spatiotemporal properties of remapping cells in detail and found that on average, these cells’ pRFs shift gradually from their cRF locations to near their fRF locations, from about 100 ms before the saccade onset to about 100 ms after the saccade offset. Importantly, the stimuli (small dots) for measuring the pRFs were flashed before the saccade onset and there were no reafferent retinal inputs to the pRFs during or after the saccade ^6^. Thus, the entire remapping time course across saccades must be driven by CDs ^3^. In addition to CD-driven forward remapping, there is also attention-driven convergent or compressive remapping of RFs toward the initial fixation and saccade target ^7,8,5,6,9^. Since convergent remapping can occur without eye movements and since forward remapping is the dominant RF dynamics around the time of saccades ^5,6^, we focus on forward remapping in this paper.

Forward remapping was recognized, at the time of its discovery, as a neural correlate of transsaccadic visual stability (TSVS), the phenomenon that the world appears continuous and stable despite saccade-induced changes of retinal images ^1^. The underlying computation, however, was unclear. Our recent measurements of the spatiotemporal properties of pRFs in LIP and FEF led to a specific circuit model that links forward remapping of RFs to transsaccadic updating of stimuli’s retinotopic positions for achieving TSVS ^6,10^. The model (Fig. 1, panels a and b) uses symmetric, center-excitation/surround-inhibition connections to store a relevant stimulus’ location as a population activity among cells tuned to different retinal locations. This memory activity bump is updated backward across each saccade as a wave by CD-gated directional connections. Gradual forward remapping of RFs is exactly equivalent to gradual backward updating of the corresponding population activities and the associated stimulus positions (under the assumption that decoders always consider cells’ activities as evidence for a stimulus in their classic RF locations) ^6,10,11^. The circuit effectively acts as an integrator of the backward motion of the activity bump, and the cumulative effect of the backward updating across the saccade is a subtraction of the saccade vector from a stimulus’ presaccadic retinotopic position to produce its correct postsaccadic retinotopic position. As such, the model fulfils the required computation that when the eye moves in one direction by a certain magnitude, the neural representation of the retinotopic position of a stimulus is updated in the opposite direction by the same magnitude ^6,10^. According to this model, transsaccadic perception is stable because the presaccadic retinal position of a stationary object in the world is updated to match the postsaccadic (reafferent) retinal position of the same object.

**Fig. 1.**
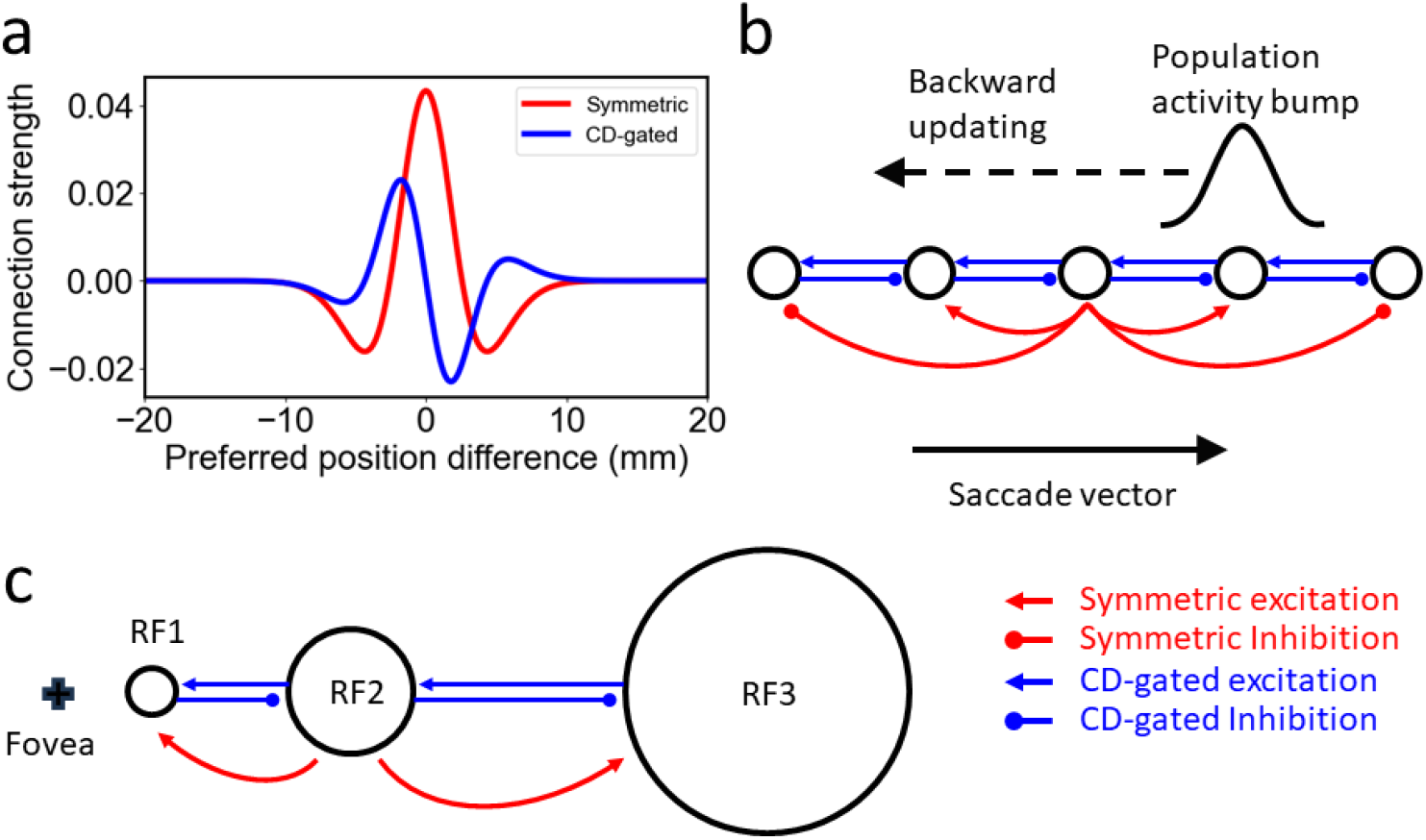
Our circuit model for RF remapping and population-activity updating across saccades. (a) Lateral connection strengths among model LIP/FEF cells as a function of the difference between the cells’ preferred retinotopic positions (RF centers). The positions can be expressed in either cortical space (mm, shown here) or visual space (deg). Symmetric, center-excitation/surround-inhibition connections (red) store a relevant stimulus’ position as a population activity bump, and directional connections (blue, shown here for rightward saccades) are gated by CDs to update the activity bump by shifting it against the saccade direction. (b) Schematic of the connections among cells whose RFs (open circles) are at different retinotopic locations. To avoid clutter, only the CD-gated connections among the nearest neighbors, and only the symmetric connections from the middle cell, are shown. During a rightward saccade (solid black arrow), the population activity bump (black curve, not drawn to the scale of the cell spacing) that represents a stimulus’ position is gradually shifted leftward (backward updating, dashed black arrow) by the CD-gated connections. Equivalently, a given cell receives lateral inputs from the cells on its right and its RF appears to gradually shift rightward (forward remapping) over time. (c) Schematic of remapping/updating among three cells of different eccentricities/RF sizes in the visual space. The cross indicates the fovea representation.

A key feature of the model is that across a saccade, a cell receives lateral inputs from other cells with different RF locations/eccentricities via the CD-gated connections among the cells. This raises the question of whether cells’ RFs transiently change sizes when they are remapped to different locations/eccentricities. Consider cells 1 and 3 with RF1 and RF3 during a rightward saccade (Fig. 1c). When cell 1 receives lateral inputs from cell 3 as the activity wave travels leftward, cell 1’s RF is remapped rightward to the location of RF3 (i.e., cell 1 appears to respond to stimuli in RF3). But RF3 has a greater eccentricity and thus larger size compared with RF1. Would cell 1’s RF appear larger when it is remapped to RF3 or would it maintain its original size of RF1? Although we raised this question before ^11^, to our knowledge, there has been no study addressing it. A previous study stated that forward remapping expands RFs of LIP cells ^4^. However, that study did not measure RF sizes and the statement was meant for the observation that during remapping, pRFs cover the midpoint between the cRFs and fRFs. More importantly, when a cell’s pRF shifts gradually from its cRF to fRF over time across a saccade ^6^, its size integrated over time can appear larger even if at any given time (or small interval of time) the size remains constant. Therefore, to address the question above, one must divide the remapping time course into small intervals, measure a cell’s RF size and eccentricity for each interval, and determine whether the RF size increases with the eccentricity.

A closely related, open question concerns the spatial uniformity of remapping/updating across the visual field. When the eye rotates in one direction by a certain visual angle, the retinal representation of the whole scene moves in the opposite direction by the same angle. This implies that the presaccadic retinal position of a stimulus should be updated by subtracting the same saccade vector *no matter where* the stimulus is in the visual space. However, when a stimulus appears at very different eccentricities, it is processed by cells with very different RF sizes and by different cell numbers per deg of visual space (cortical magnification factor). How does a highly nonuniform neural representation of the visual space ensure uniform transsaccadic updating across the visual space?

Our previous circuit model used the simplifying assumption that the cells represent the visual space uniformly (Fig. 1b). However, to address the above two questions, we must incorporate the fact that the RF size and cortical magnification factor changes drastically with eccentricity (Fig. 1c). In this study, we introduced eccentricity dependence into our model and mathematically analyzed the model behavior. We found that surprisingly, when the CD-gated connections are chosen to achieve uniform updating across the visual space (but not the cortical space), the model predicts no change to a cell’s RF size during remapping despite its receiving lateral inputs originated from other cells with different eccentricities/RF sizes. In contrast, if the updating is uniform across the cortical space (but not the visual space), a cell’s RF changes size according to the eccentricity of the remapped location at a given time. Between these two cases, the remapping is nonuniform in either space, and a cell’s RF size changes by various degrees during remapping. Finally, by analyzing our previous single-unit data from LIP and FEF, we showed that remapping magnitudes for cells with large and small eccentricities are not significantly different from each other, and confirmed the prediction that the RF size does not change significantly with the eccentricity during remapping. Our work thus suggests the first circuit mechanism for achieving uniform remapping/updating as required by transsaccadic perceptual stability across the entire visual space.

## Results

### A circuit model for uniform remapping/updating in the visual space

For a given saccade, the forward remapping and backward updating are along the axis that contains the saccade direction. It is thus sufficient to model just one spatial dimension which we call horizontal; saccades along other axes can be treated similarly. The details of the circuit model can be found in Methods and Supplemental Information. To facilitate understanding, we use dual representations of the same circuit model: one in the cortical space (measured by distance *x* along the cortical surface) and the other in the visual space (measured by degree *y* of visual angles). It is known that for the primary visual cortex the mapping from *x* to *y* is approximately exponential ^12^. We assume a similar relationship for LIP/FEF:

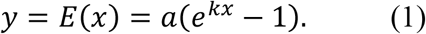

Note that y = 0 when x = 0, and we can use 0 position to represent the fovea/fixation in both spaces. *x* and y also represent the eccentricity in these spaces. The parameters a > 0, k > 0 are constant for a given cortical area. From Equation 1, we have:

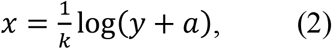

the logarithmic compression of the visual space to the cortical space, and the cortical magnification factor:

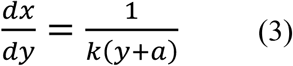

that decrease with eccentricity y. Since cell density over the cortical space (*dn/dx* where *n* is the cell number) is uniform (constant over *x*) ^13^, Equation 3 implies that the cell density over the visual space (*dn/dy*) decreases with eccentricity *y* of the visual space:

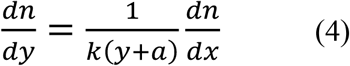

The size of a cell’s classic RF, measured in the visual space (e.g., on a display screen), is proportional to the eccentricity *y* of the cell’s RF (as we show in Fig. 8 with our LIP/FEF data):

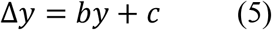

where the parameters *b* > 0, *c* > 0 are also constant for a given cortical area, and *c* is the RF size at fovea. Our circuit model shows the same relationship when the symmetric, center/surround connectivity is uniform over the cortical space. We show in Supplemental Information that at the stable state, a cell at *x* has an RF size of 2*d*, covering [*x-d, x+d*] in the cortical space, where *d* is determined by the attractor dynamics of the symmetric connections and independent of *x*. Using Equation 1, the corresponding RF size Δ*y* in the visual space at eccentricity *y* is:

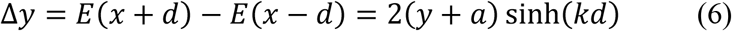

in agreement with Equation 5, with *b* = 2 sinh(*kd*) and *c* = 2*a* sinh(*kd*). Additionally, for two cells with eccentricities *y*_1_ and *y*_2_ that have abutting but non-overlapping RFs in the visual space, their separation in the cortical space is always 2*d* independent of the eccentricities (this separation is 2 mm in the primary visual cortex ^13^).

The CD-gated, antisymmetric connectivity pattern is chosen to shift the population activity bump backward across saccades while maintaining the shape of the bump in the cortical space ^6,10,14^. This connectivity can be scaled by an amplitude (*J*) which, after divided by the neuronal time constant τ, determines the remapping/updating speed in the cortical space ^14^ (see Methods):

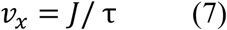

(the remapping and updating speeds have equal magnitude but opposite signs ^6,10,11^). In our previous simulations, *J* depends on only the CD strength as a function of time, *J* = *g*_*cd*_(*t*), chosen to make the cumulative remapping/updating magnitude equal the saccade magnitude (for stimuli appeared well before the saccade ^6,10^). Since we now consider a nonlinear function between the cortical space and visual space (Equation 1), the same activity bump has different speeds in the two spaces, and we need to choose *J* to achieve the desired uniform remapping/updating across the visual space *y*. By differentiating Equation 1 with respect to time, we obtain the relationship between the speeds in the two spaces:

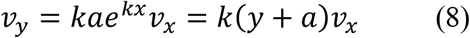

where *v*_*x*_ = *dx*/*dt* and *v*_*y*_ = *dy*/*dt* are the speeds in the cortical and visual spaces, respectively.

We first consider two special cases. (1) If we let *J* = *g*_*cd*_(*t*) as we did previously, then *v*_*x*_ (Equation 7) is uniform across the cortical space (constant over *x*, not time), but *v*_*y*_ (Equation 8) increases with eccentricity *y* in the visual space. (2) If, instead, we let

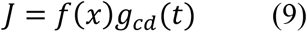

where:

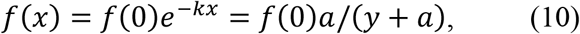

then *v*_*y*_ is uniform over the visual space (constant over *y*, not time), but *v*_*x*_ decreases with *x*. Since the circuit is effectively an integrator of the remapping/updating speed to produce a total, accumulative shift across saccades, (non)uniform speed in a space implies (non)uniform remapping/updating magnitude in that space. Thus, cases 1 and 2 specify the connectivity in the circuit model for realizing uniform remapping/updating in the cortical space and visual space, respectively.

We next investigated, for the above two cases, how the model predicts RF-size change during remapping. Consider a cell at eccentricity *y*_1_ with RF size Δ*y*_1_ that is remapped to eccentricity *y*_2_ with RF size Δ*y*_2_ at a given time relative to the saccade onset. We found that in case 1, the model predicts:

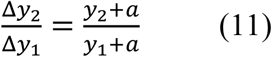

whereas in case 2, the prediction is:

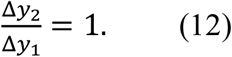

That is, when the remapping/updating is uniform in the cortical space but not visual space, the cell’s RF size changes with the eccentricity during the remapping (Equation 11), but surprisingly, when the remapping/updating is uniform in the visual space but not cortical space, the cell’s RF size does not change during the remapping (Equation 12) despite its receiving lateral inputs from other cells with different eccentricities/RF sizes.

Fig. 2 demonstrates Equation 11. Consider cells 1 and 2 whose retinotopic locations are *x*_1_ and *x*_2_ in the cortical space, and *y*_1_ and *y*_2_ in the corresponding visual space (vertical red ticks in Fig. 2). Each curve in the figure represents the population activity bump evoked by a retinal stimulus at the arrow of the same color and style (the peak activity is aligned with the corresponding arrow). As explained below, the dashed curves can also represent activity bumps updated from the solid curves at a fixed time relative to the saccade onset. For each cell in either space, the stimuli at the pair of green and blue arrows evoke population activity bumps that just include the cell (red tick), so the region between the green and blue arrows is the cell’s RF size. During a rightward saccade, cell 1 receives lateral inputs originated from cell 2. Because *v*_*x*_ does not depend on *x*, the population activity bumps evoked by stimulating any part of cell 2’s RF will travel toward cell 1 at the same speed in the cortical space. In particular, the solid green and blue bumps evoked by the solid green and blue arrows will shift to the dashed green and blue bumps (Fig. 2a) at the *same* time relative to the saccade onset. Thus, when cell 1 receives maximal inputs from cell 2 so that cell 1’s RF is remapped to the location of cell 2, cell 1’s RF size equals that of cell 2 in the cortical space. Since cell 2 has a greater eccentricity and thus a larger RF than cell 1 in the visual space (Fig. 2b), cell 1’s RF becomes the larger RF of cell 2. We conclude that when remapping/updating is uniform in the cortical space, a cell’s RF size in the visual space will change according to the eccentricity of its remapped RF location at a given time relative to the saccade onset (Equation 11).

**Fig. 2.**
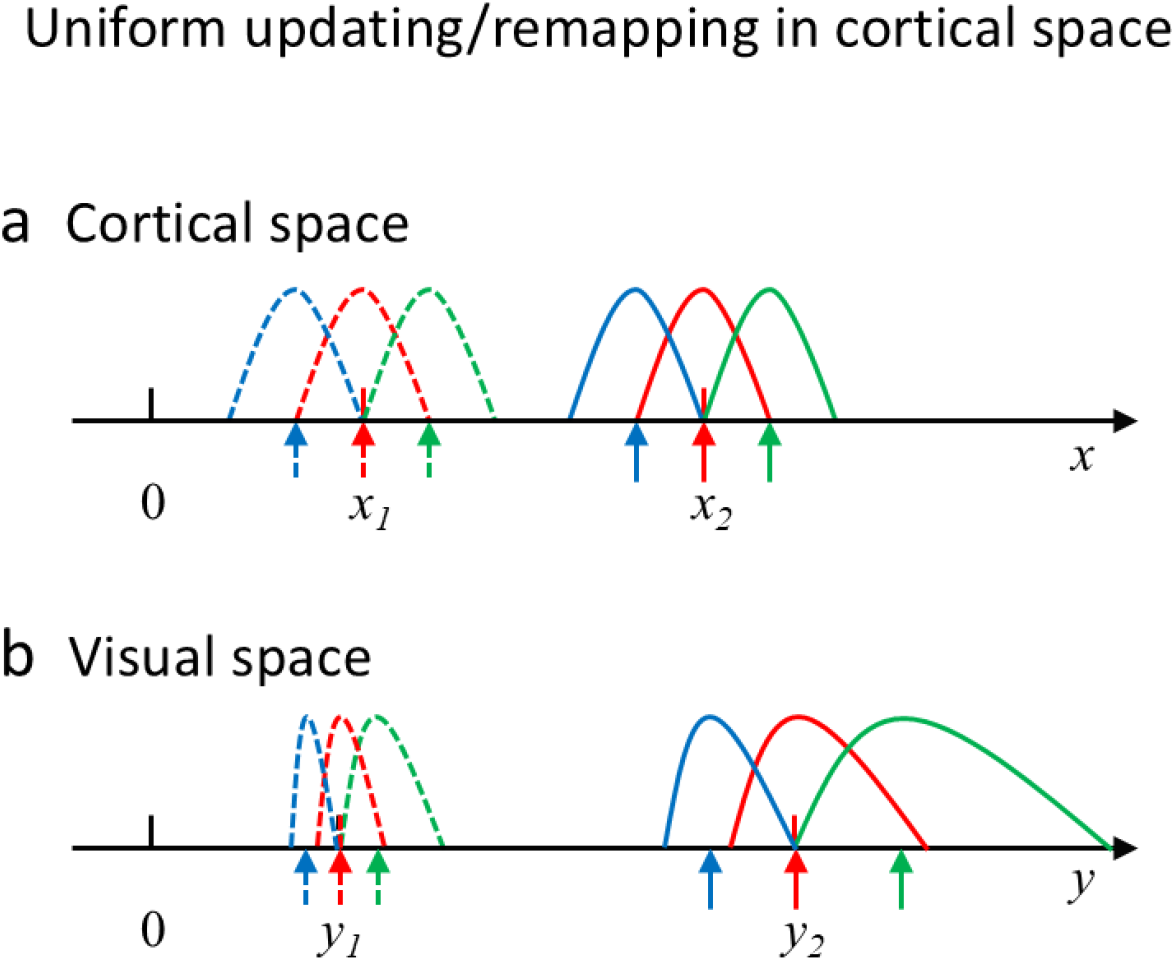
Uniform updating/remapping in the cortical space but not visual space. Consider two cells, indexed 1 and 2, whose retinotopic locations are *x*_1_ and *x*_2_ in the cortical space, and *y*_1_ and *y*_2_ in the corresponding visual space (short vertical red ticks). Each curve in the figure represents the population activity bump evoked by a retinal stimulus at the arrow of the same color and style (the peak activity is aligned with the corresponding arrow). During a saccade, cell 1 receives lateral inputs from cell 2 via CD-gated connections, so the dashed curves also represent population activity bumps updated from the solid curves at a fixed time relative to the saccade onset. Whereas activity bumps in the cortical space maintain the same size and shape, they change with eccentricity in the visual space (see text). (a) Cortical space. Since the speed of updating/remapping is uniform in this space, at a given time, the spacings among the updated activity bumps (dashed curves) are identical to those among the original activity bumps (solid curves). (b) Visual space. The speed of updating/remapping is not uniform in this space, with the bump at smaller eccentricity (solid blue curve) moves slower than that at larger eccentricity (solid green curve). Consequently, the updated activity bumps (dashed curves) become closer to each other.

Figs. 3 and 4 provide an intuitive explanation of Equation 12 and the details can be found in Supplemental Information. In Fig. 3, the solid green and blue arrows around cell 2 are at the positions identical to those in Fig. 2; they evoke the solid blue and green activity bumps that just include cell 2 and thus indicate the right and left borders of cell 2’s RF. However, the updating of the activity bumps are different from that in Fig. 2. Because the updating speed is now uniform in the visual space, at a given time relative to the saccade onset, the spacings between the three updated activity bumps (dashed curves in Fig. 3b) must equal the spacings of the original activity bumps (solid curves in Fig. 3b). This means that at time T (relative to the saccade onset) when the dashed red curve shows maximal activation of cell 1, the dashed green and blue curves show no activity at cell 1. Thus, the solid green and blue arrows are outside cell 1’s remapped RF at time T. Fig. 4b shows that the solid green and blue arrows around cell 2 must cover a size equal to cell 1’s RF to activate cell 1 at time T (Equation 12).

**Fig. 3.**
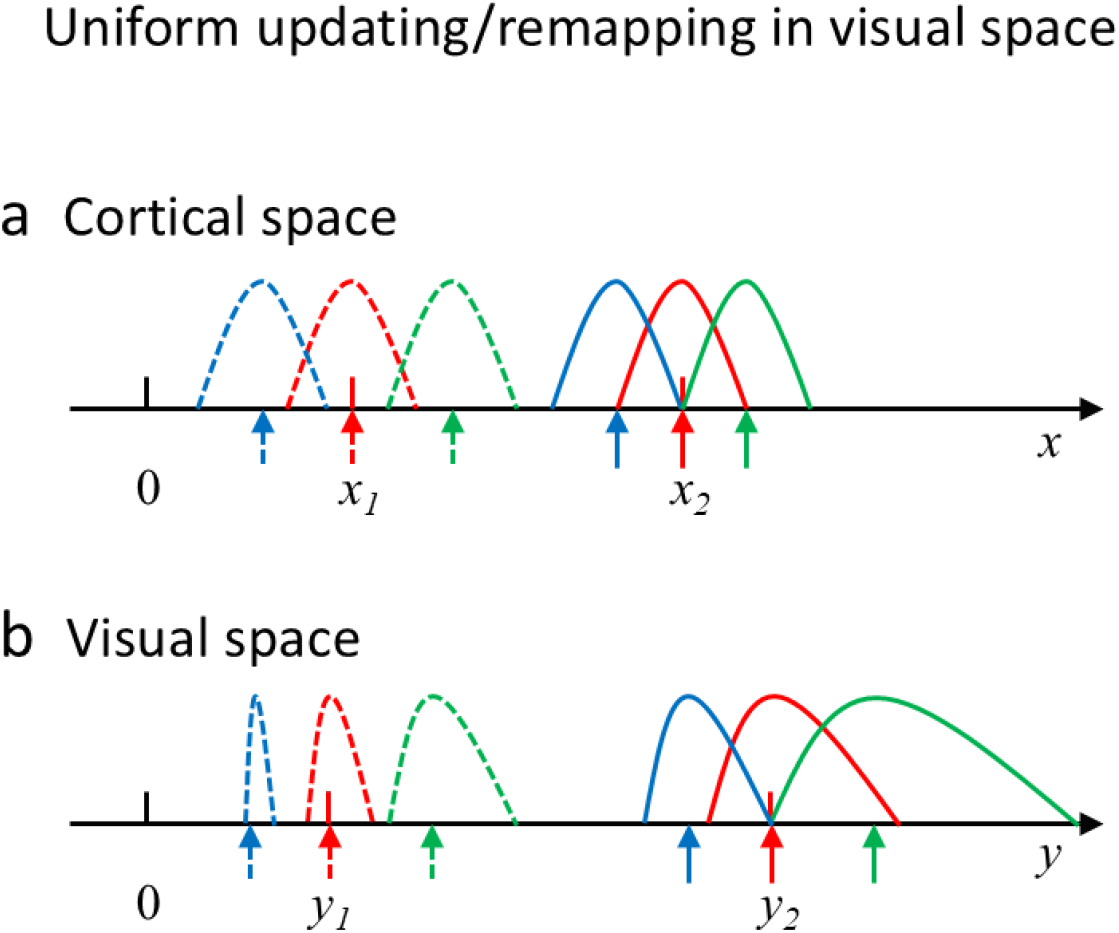
Uniform updating/remapping in the visual space but not cortical space. The format is identical to that of Fig. 2. (a) Cortical space. The speed of updating/remapping is not uniform in this space, with the activity bump at smaller eccentricity (solid blue curve) moves faster than that at larger eccentricity (solid green curve). Consequently, the updated activity bumps (dashed curves) become farther apart to each other. (b) Visual space. Since the speed of updating/remapping is uniform in this space, at a given time, the distances between the peaks (arrows) of the updated activity bumps (dashed curves) are identical to the corresponding distances between the original activity bumps (solid curves).

**Fig. 4.**
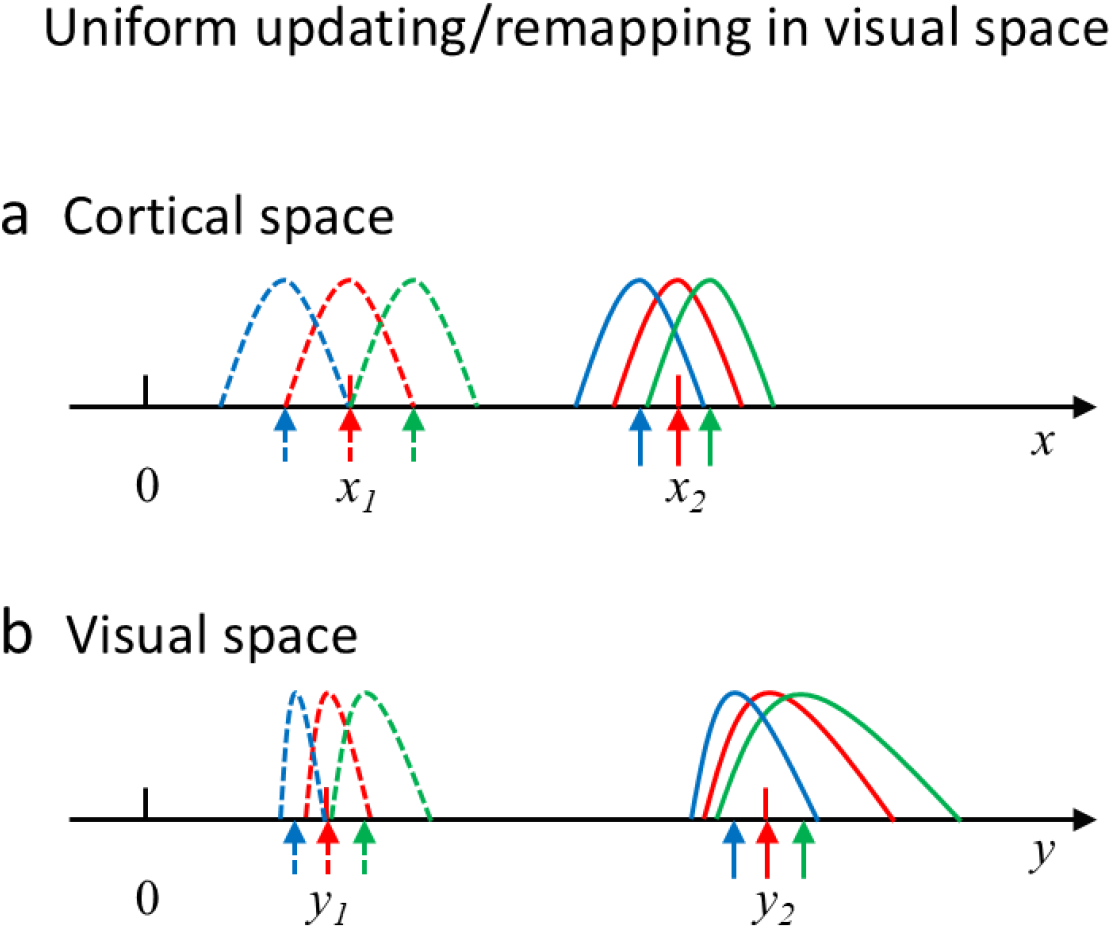
Uniform updating/remapping in the visual space but not cortical space. The format is identical to that of Figs. 2 and 3. The figure is identical to Fig. 3 except that here the dashed blue and green arrows indicate cell 1’s RF borders; that is, the population activity bumps (dashed blue and green curves) just include cell 1 at the red tick. The figure explains why RF sizes do not change during remapping/updating that is uniform in the visual space.

We finally consider the general case where:

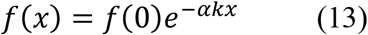

where 0 ≤ α ≤1. Note that α = 0 and 1 correspond to the two special cases considered above, respectively; other values of α cover intermediate cases where the speed of remapping/updating is not uniform in either the cortical space or visual space. In Supplemental Information, we show that when a cell with RF eccentricity *y*_1_ is remapped to eccentricity *y*_2_, its RF size at that time changes from Δ*y*_1_ to Δ*y*_2_ according to

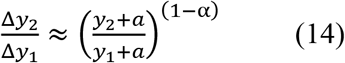

The approximate expression reduces to the exact results for the two special cases of α = 0 and 1 in Equations 11 and 12.

We have simulated our circuit model to confirm the above analysis. Examples for the two special cases are shown in Fig. 5. When the speed of remapping/updating is uniform in the cortical space (Fig. 5a, blue) but not in the visual space (Fig. 5a, red), the RFs changes size during remapping according to eccentricity (Fig. 5b). By contrast, when the speed of remapping/updating is uniform in the visual space (Fig. 5d, red) but not in the cortical space (Fig. 5d, blue), the RFs do not change size during remapping (Fig. 5e). In both cases, the population activity bumps always change sizes during updating (Fig. 5, c and f). The figure also illustrates that tuning curves (RFs are spatial tuning curves) and the corresponding population activities generally do not have the same shape ^11,15–18^.

**Fig. 5.**
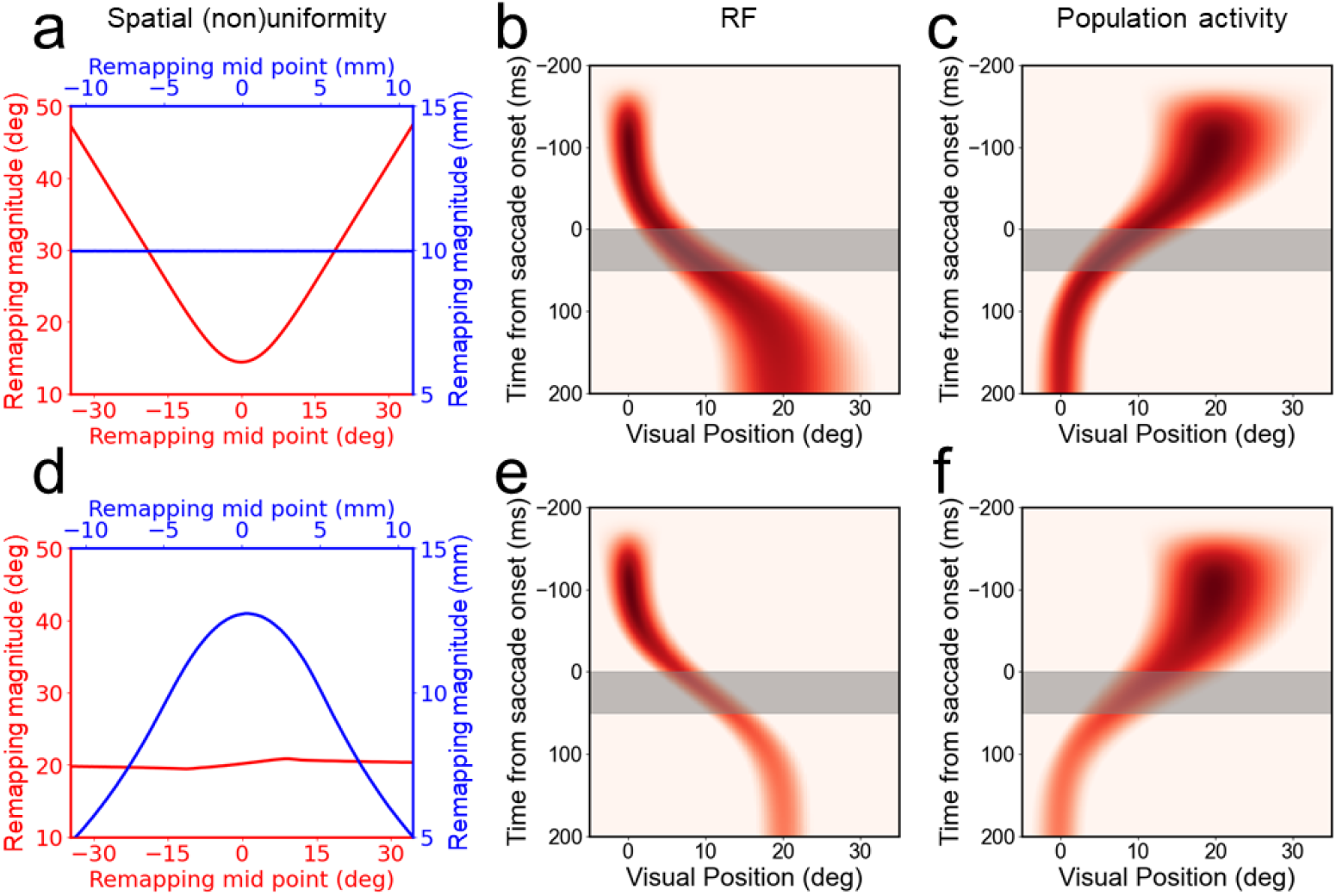
Simulations of forward RF remapping and backward population-activity updating across a 20-deg rightward saccade with the circuit model. The top row shows case 1 in which remapping/updating is uniform in the cortical space but not visual space. The bottom row shows case 2 in which remapping/updating is uniform in the visual space but not cortical space. The foveal position in both spaces is 0. The left column (a, d) shows total remapping magnitude as a function of the mid-point position. The red and blue colors indicate results for the visual space and cortical space, respectively. The middle column (b, e) shows the heatmap of a cell’s pRF position in the visual space as a function of time relative to the saccade onset. The gray-shaded areas indicate the saccade duration. Stimuli for measuring the pRF is flashed 200 ms before the saccade onset. The right column (c, f) shows the heatmap of all cell’s population response, evoked by a flash at 20 deg in the visual space and 200 ms before the saccade onset, as a function of time. During remapping, the pRF size changes with eccentricity in case 1 (panel b) but remains constant in case 2 (panel e). In both cases, the population activity changes size with eccentricity (panels c and f).

### Analyses of LIP and FEF single-unit data

To test the theory above, we re-analyzed our previously published single-unit data from LIP and FEF ^6,10^ to check whether or not the remapping is uniform over the visual space, and whether or not the RFs change sizes during remapping. For the cells in each area that passed the screening procedure ^6,10^ (Methods), we divided them into the small-and large-eccentricity groups according the medium eccentricity of the cells. Since the remapping magnitude accumulates across a saccade and ends about 100 ms after the saccade offset, or 150 ms after the saccade onset, we measured the final remapping magnitude of each cell in the time window [100, 150] ms after the saccade onset and normalize it by the saccade amplitude used for recording the cell (so that we can pool cells recorded with different saccade amplitudes). The results are shown in Fig. 6. The left panels show that the means and standard errors of the eccentricities of the two groups of cells; their means are significantly different as expected (two-sided Wilcoxon rank-sum test; p = 9.7×10^-10^ for FEF; p = 1.4×10^-8^ for LIP). The right panels show that their remapping magnitudes are similar and not significantly different (p = 0.86 for FEF; p = 0.81 for LIP). This provides the first evidence for the desired uniform remapping/updating over the visual space in LIP and FEF.

**Fig. 6.**
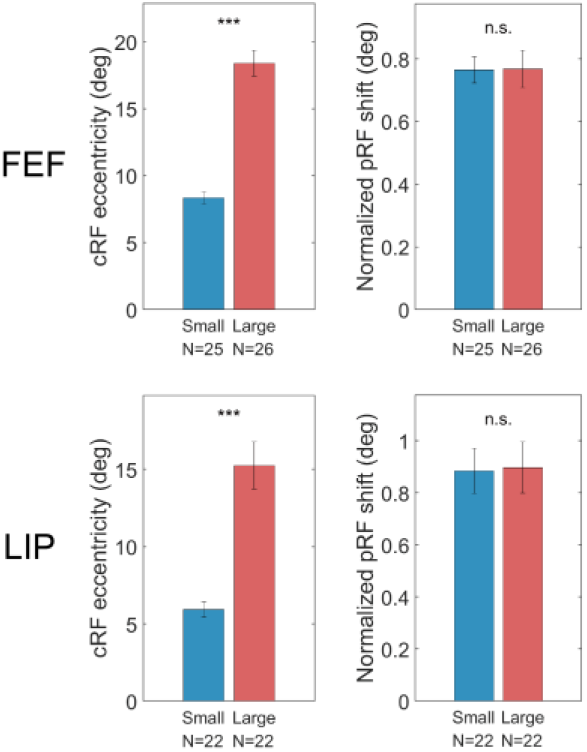
Spatially uniform remapping in LIP and FEF. The left panels show that the small- and large-eccentricity groups of cells have significantly different mean eccentricities. The right panels show that the two groups have similar pRF remapping magnitudes.

With the evidence for spatially uniform remapping in Fig. 6, our analysis and simulations predict that when a given cell’s RF shifts to different eccentricities during remapping, its size does not change with the eccentricity. We tested this prediction. We first divided a cell’s responses into four non-overlapping 50-ms time bins centered at -25 ms, 25 ms, 75 ms, and 125 ms relative to the saccade onset. For each cell and time bin, we determined the pRF size and eccentricity (Methods). Finally, we plot the pRF size change as a function of the eccentricity change. The changes are relative to the corresponding cRF size and eccentricity (during fixation without remapping). It is important to focus on these changes which shows whether the *same* cell changes its RF size when its eccentricity changes during remapping. Otherwise, the plot would be confounded by the well-known dependence of RF size on eccentricity among *different* cells. Fig. 7 shows the results for FEF and LIP. During the remapping, the RF-size change does not vary significantly with the RF-eccentricity change in each area (linear mixed model, slopes and p values in the figure), consistent with our prediction. To demonstrate the robustness of the results, in Supplemental Information, we show results similar to Figs. 6 and 7 when we changed the parameter for defining RF borders (contour criterion; Methods) from 0.6 to 0.7.

**Fig. 7.**
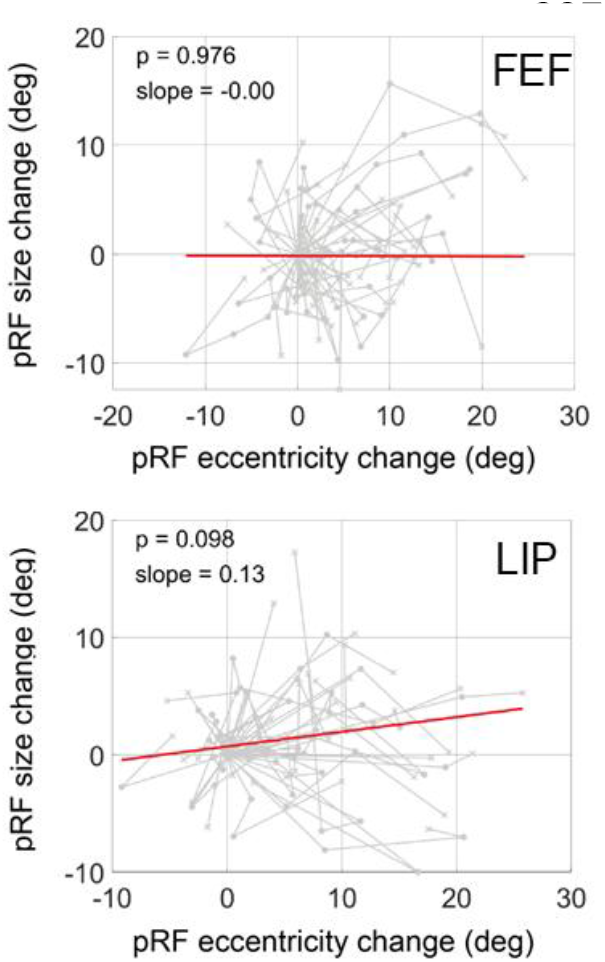
The change of pRF-size as a function of the change of eccentricity during remapping for cells in FEF (top) and LIP (bottom). For each cell, the changes are relative to its cRF size and eccentricity during fixation (without remapping). The same cell’s results in different time bins (see text) are linked by gray lines. The cells numbers are the same as in Fig. 6. The red line in each plot is the fit of linear mixed model, which we use because different data points of the same cell are not independent samples.

A potential problem with the data analysis in Fig. 7 is that because we had to divide the responses of small numbers of cells into time bins of 50 ms, there may not be enough spikes in each bin to show a reliable dependence of pRF size on eccentricity during remapping. To control for this possibility, we analyzed the cRF data from the initial fixation period (without remapping) and used the same 50-ms bin size for counting spikes. We found that the cRF-size as a function of eccentricity among different cells is still significant over each of the 50-ms bins even when we matched the pRF cell numbers (Fig. 8). Therefore, the insignificant change of the pRF size during remapping in Fig. 7 is not because of the small bin size or small numbers of cells.

**Fig. 8.**
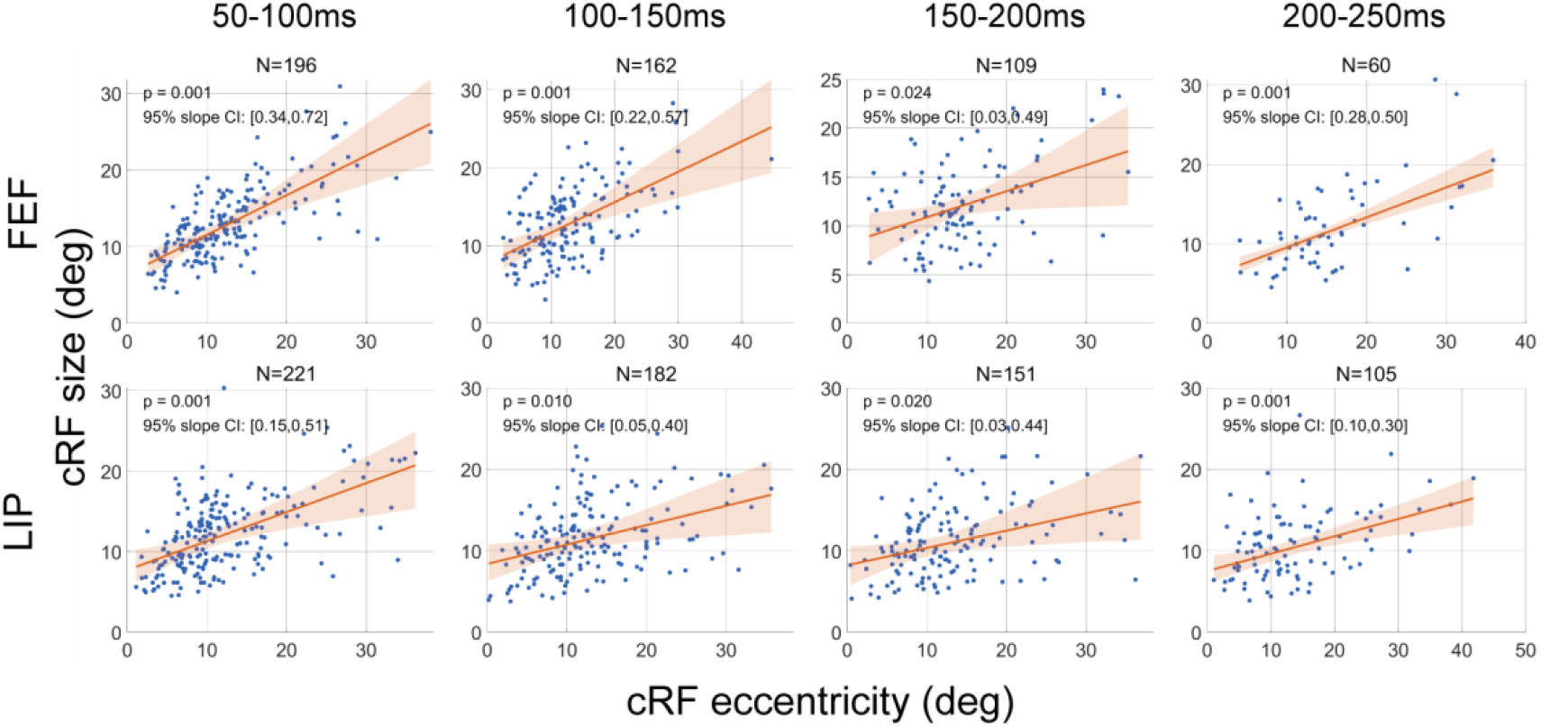
Dependence of cRF size on eccentricity in FEF and LIP. This is a control analysis of the FEF and LIP data to show that the lack of significance in Fig. 7 is not because of the small size of time bins and small numbers of cells. We analyzed cRF sizes in four, 50-ms time bins (top, times relative to the onset of the probe stimuli). The blue dots show the cells with significant responses. We then randomly sampled a subset of the cells to match the cell numbers in Fig. 7, and estimated the mean (red lines) and 95% confidence intervals (shaded areas) from 1000 such subsamples. The slopes of the red lines are significantly different from 0 in all cases.

## Discussions

We addressed two closely related issues in this paper. First, perisaccadic RF remapping can transiently change a cell’s eccentricity by a large amount. Since RF sizes depend on eccentricities, does a given cell change its RF size during remapping? In our circuit model, remapping occurs because cells receive lateral inputs originated from other cells. This raises a similar question of whether or not a cell changes its RF size when it receives inputs from other cells with very different eccentricities and RF sizes? Second, for forward remapping, or equivalently backward updating, to be a useful mechanism for achieving TSVS, it must be uniform across the visual space: for a given saccade, the updating of a stimulus’ retinotopic position should be a subtraction of the saccade vector regardless of where the stimulus appears in the visual space. How is this desired uniform remapping/updating realized based on cells with very different RF sizes and cortical magnification factors at different eccentricities of the visual space? To address these questions, we introduced the standard nonlinear relationship between the cortical and visual spaces into our circuit model. Our analysis and simulations demonstrate that the CD-gated connections in the model can be chosen to achieve uniform remapping/updating in the visual space, and in this case, a cell’s RF size does not change during remapping despite its receiving inputs from other cells with different eccentricities/RF sizes. By contrast, if the connections are chosen such that remapping/updating is uniform in the cortical space (but not in visual space), a cell’s RF size does change during remapping. We analyzed our previously published data from LIP and FEF and found that remapping magnitudes (normalized by the saccade magnitudes) are similar for cells of large and small eccentricities. This is the first physiological evidence of uniform remapping in the visual space. We then confirmed the prediction that RF sizes did not change significantly with eccentricity during remapping. The work provides a specific circuit mechanism for realizing uniform remapping/updating and transsaccadic perceptual stability across the entire visual field.

In addition to the prediction on RF sizes during remapping, another prediction of the model is Equation 10 which specifies the decrease of the CD-gated connection magnitude with eccentricity to ensure uniform remapping/updating in the visual space. Additionally, we predict that the translational component of perisaccadic mislocalization (the dominant component in the dark ^19^) must be uniform across the visual space. This is because according to our model, translational mislocalization is produced by the temporal relationship between the flash time of a stimulus and the CD time course ^10^ and this relationship does not change with the stimulus location when remapping/updating is uniform over the visual space.

Most physiological studies of perisaccadic remapping and our circuit model for remapping/updating focus on stimulus *position*. What, then, happens to stimulus *features/contents* across saccades? A small number of physiological studies that examined this question found different degrees of feature remapping in different brain areas with different methods ^20–22^. By contrast, many psychophysical experiments examined feature, form, object, or even scene perception across saccades. A popular paradigm is saccadic change blindness: a stimulus is presented before a saccade and then changed during the saccade ^23^. Because of saccadic suppression, subjects usually do not see the change when it happens but have to report after the saccade whether or not they noticed a change. Typical findings are that subjects often fail to detect large changes, similar to the results of fixational change blindness (in which a blank screen, instead of a saccade, separates a pair of stimuli to prevent a direct, temporal comparison). Such findings suggest that only a small number of attended image patches are represented under fixation and get remapped/updated across saccades, and even for the remapped/updated image patches, not all features/details in them are stored and carried over saccades. Further studies are needed to determine the properties of feature, form, object, and scene remapping/updating across saccades, their relationship to attention and perception, and their dependence on tasks.

As noted in Introduction, according to our model, a stationary object in the world appears stable across a saccade because its presaccadic retinotopic position is updated to its correct postsaccadic retinotopic position (by the subtraction of the saccade vector), and the updated position agrees with the new (reafferent) retinal inputs from the same object after the saccade ^10^. A mismatch between the updated position and the reafferent position may drive learning/plasticity to adjust the CD-gated connections and correct the mismatch. Whether and how this learning process operates under developmental and pathological conditions are open questions. A related question is whether there is a threshold for the mismatch to reach before the learning is triggered. Indeed, the paradigm of saccadic change blindness has also been used to investigate subjects’ sensitivity to changes of stimulus positions, and the conditions for both low and high sensitivities have been reported ^24,25^. A further question is whether the sensitivity topositional changes decreases, and the learning threshold increases, with eccentricity, given that visual acuity decreases with eccentricity. If so, remapping/updating would not be perfectly uniform across the visual space, and the circuit mechanism we proposed here, like all models, must be an idealized approximation of biological reality.

## Methods

### Circuit model

We modeled a one-dimensional array of retinotopic LIP/FEF units covering the horizontal axis. We use a dual representation of the same circuit model: in the cortical space measured by distance *x* in mm along the cortical surface and in visual space measured by *y* in degrees of visual angles. We first determined the model in the cortical space, and then used Equation 1 in the main text to specify the relationship between the cortical space and the visual space. In the cortical space, each unit is governed by the equations:

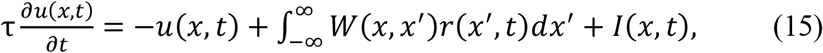

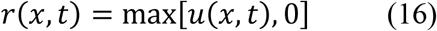

where *u*(*x, t*) and *r*(*x, t*) represent, respectively, the membrane potential and firing rate of the unit at location and time (*x, t*), *τ* is the membrane time constant, *W*(*x, x*^′^) is the recurrent (lateral) connection strength from neuron at *x*^′^ to neuron at *x*, and *I* is the feedforward inputs to LIP/FEF which originate from the retina. *W*(*x, x*^′^) is a sum of two parts: : (1) symmetric, center-excitation/surround-inhibition connections modeled as a weighted difference between two Gaussians: *W*_*sym*_ = *J*_*exc*_*G*(*x, x*^′^, σ_*exc*_) − *J*_*inh*_*G*(*x, x*^′^, σ_*ing*_) where 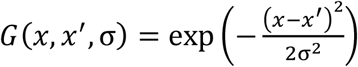, and (2) directional connections gated by the saccade CD, with predominantly excitation and inhibition in the backward and forward directions of the saccade, respectively. For the simulations in this paper, we let *J*_*exc*_ = 0.11, σ_*exc*_ = 2 mm, *J*_*inh*_ = 0.06, σ_*inh*_ = 3.19 mm. We modeled the CD-gated connections as the antisymmetric, spatial derivative of the symmetric connections: 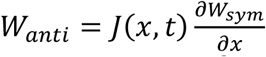 where *J*(*x, t*) = *f*(*x*)*g*_*CD*_ (*t*) with the CD gating factor 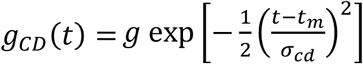. This choice of *W*_*anti*_ is known to preserve the shape of the activity bump when the spatial factor *f*(*x*) is a constant ^14^ and our simulations show that the shape is well maintained even when *f*(*x*) is not a constant. We set *t*_*m*_ to be the mid time of the saccade duration assumed to be 50 ms, and σ_*cd*_ = 60 ms. The peak CD *g* depends on the saccade and for convenience, we merged it with *f*(*x*). Other forms of *g*_*CD*_(*t*) work too ^6,10^. As explained in the text, we considered cases where *f*(*x*) is a constant and where *f*(*x*) varied with *x* according to Equation 11. In the former case we let *f*(*x*) = 1.36, and in the latter case we let *f*(*x*) = 2.65*e*^−0.125*x*^. This means *k* = 0.125/mm in Equation 1, and we set *a* = 8.05° in that equation for our simulations. These parameters map 20 mm of the cortical space to 90° of the visual space. The blue curve of Fig. 1a shows only the 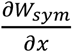 part of the directional connections. The speeds of remapping and updating have opposite signs, and their absolute value in the cortical space is given by ^14^:

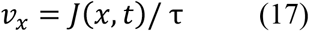

For visual inputs, we considered flashed spots on retina as in the actual experiment of Wang et al.^6,10^ A spot flashed on retina is filtered both spatially and temporally when it reaches LIP/FEF so we modeled its input to LIP/FEF units as a spatial Gaussian function and a temporal gamma function:

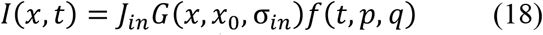

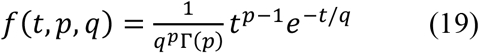

where *x*_0_ is the retinotopic position of the flash, and *p* and q are the shape and scale parameters, respectively. We let σ_*in*_ = 1.5 mm, *J*_*in*_ = 4, *p* = 6, *q* = 8 ms so the delay from the retinal flash to the peak of the LIP/FEF input is (*p*-1)*q* = 40 ms.

### Data analysis

We reanalyzed our LIP and FEF single-unit data reported in our previous publications ^6,10^ to test two predictions: (i) for the pRFs we measured with flashes before the saccades, the final magnitude of forward remapping (which completes about 150 ms after saccade onset) should be independent of the cRF eccentricity; and (ii) when the pRFs shift in space during remapping, their sizes at a given time (or small time interval) should remain unchanged. Detailed information regarding the experimental design, data collection, and data analysis can be found in those publications. Briefly, single-unit activity was recorded from the LIP and FEF of monkeys while they performed a delayed saccade task. For each unit, RFs were measured across four distinct time periods— current (presaccadic before the target onset), delay (presaccadic after the target onset), perisaccadic, and future (postsaccadic)—by presenting a probe stimulus at one of multiple array locations during each period of every trial. These RFs are hereafter referred to as the cell’s cRF, dRF, pRF, and fRF, respectively. For the current purpose, we focused on the cells’ pRFs measured during the perisaccadic period when the remapping direction is largely forward. We modified and extended our previous analysis method to include the calculation of the RF size. We fully describe the method below including some details inadvertently left out in our previous publications. The Matlab code for the data analysis and the Python code for the model simulations will be posted online after the publication. The data analysis code also reproduces the results in our previous publications.

#### Stimulus presentation time

When the computer code for stimulus presentation is executed, it takes a variable delay, in the range of tens of ms, for the stimulus to actually appear on the display screen. To determine the stimulus presentation time more accurately, we attached a photocell at the lower-right area of the screen that was turned on and off with the stimulus in the code (the area was covered), and recorded the photocell output via a Labjack data acquisition unit in each period of each trial. We used the onset and offset times of the output as a proxy for the stimulus onset and offset times. For the perisaccadic period, it is important that all probe stimuli for measuring the pRF appeared before the saccade onset to ensure that they were all in the same retinotopic reference frame. We therefore excluded trials in which the stimulus offset time was later than the saccade onset time. The resulting data set was published previously ^10^ and used in the current study. The saccade onset time was the time when the eye speed first exceeded 30 deg/sec ^26^. The eye position was recorded with subconjunctival search coils sampled at 2.7KHz ^6^.

#### Significant response screening

We selected cells exhibiting significant visual responses independently for each time period and alignment condition. For each period, repeated trials were aligned to the onset of the probe stimulus; for the perisaccadic period, trials were additionally aligned to the onset of the saccade. For the screening purpose, the response was defined as the mean firing rate recorded between 50 and 150 ms after each probe onset, or between 0 and 100 ms after saccade onset (these time windows were selected to capture the majority of evoked neural activities). The baseline firing rate was defined as the mean firing rate recorded between 0 and 50 ms before the corresponding probe onset. The probe position that elicited the maximal response was identified, and a two-sided Wilcoxon rank-sum test was performed to compare responses at this position with the corresponding baseline (*α* = 0.05). Only cells that passed this test were retained for subsequent analyses. For each selected cell, spatial responses were normalized across probe positions as (*r*_*k*_ − *r*_*min*_)/(*r*_*max*_ − *r*_*min*_), where *r*_*k*_is the response at position k and *r*_*max*_ and *r*_*min*_ are the maximum and minimum responses across positions. Subtracting *r*_*min*_ reduces the influence of non-visual activity (e.g., saccade-related responses). Additionally, because the saccade target was consistently placed outside the cell’s receptive field, any contribution of saccade-related activity to the measured visual responses was minimized.

#### Well-measured RF screening

We selected cells with well-measured RFs. Following significant response screening, normalized responses across sampled probe positions were linearly interpolated and resampled on a two-dimensional grid (grid spacing = 0.1 deg along both axes) to generate an RF heatmap. A contour corresponding to a specified percentage of the cell’s maximum response (the contour criterion) was then traced. We additionally required that every probe position falling within this contour had at least five trials, and that at least 80% percent of the contour is within the grid region (the completeness criterion). If part of a contour was not within the grid region, then the relevant grid boundary is used as the contour. We used the region within the contour to calculate both the RF area and the RF center. The RF area was the number of resampled grid points within the contour multiplied by the grid-spacing squared. The RF size was defined as the square root of the RF area. The RF center was the center of mass within the contour weighted by the responses.

We previously only needed to determine the RF center location ^6,10^ whereas in the current study, we additionally needed to determine the RF size. For the RF center calculation, it is sufficient to focus on a small area around the peak response point, andwe previously used a standard value of 0.85 for the contour criterion (and showed similar results when it was set to 0.75). By contrast, for the RF size calculation, it is more reliable to include a larger area around the peak response point. However, when the contour criterion has a smaller value to include a larger area, fewer cells satisfy the completeness criterion and the results become noisier. To balance these considerations, in the present study, we used a standard value of 0.6 for the contour criterion, and showed similar results when it was set to 0.7. We also used a larger contour criterion for measuring the RF center and a smaller contour criterion for measuring the RF size. Since the results are very similar to those with a single contour criterion, we do not report them here.

#### Significant remapping screening

We selected cells showing significant pRF shifts. For each cell, the displacement of the pRF centers relative to the cRF center was computed (in degrees of visual angle), and the significance of these displacements was assessed via bootstrap simulation. Specifically, for each probe position of a given cell, per-trial spike counts were modeled as a Poisson distribution with a mean equal to the observed mean spike count. One thousand repeated simulations were then generated, with spike counts sampled from these distributions (matching the number of trials in the original experiment). To test whether the pRF center of a cell differed from the cRF center, the 1,000 bootstrapped pRF and cRF centers were projected onto the axis connecting their respective means. The fraction of overlap between these two one-dimensional projection distributions was quantified, and a shift was deemed significant if this overlap was < 5%.

We used the cells that passed the above screening steps for each period and brain area for further analyses below.

#### Forward remapping magnitudes for small- and large-eccentricity groups of cells

As we noted in the text, for the forward remapping to be a useful mechanism for TSVS, the remapping magnitude must be uniform across eccentricity in the visual space. We calculated each cell’s eccentricity as the distance between its cRF center and the fixation. The cRF center was determined with the responses 50–150 ms after the probe onset. We then divided the cells in each brain area into small- and large-eccentricity groups using the median eccentricity. Since forward remapping ends around 150 ms after the saccade onset (or 100 ms after the saccade offset), we calculated the final pRF remapping magnitude using the responses 100-150 ms after the saccade onset. The remapping magnitude is the distance between a cell’s pRF center and cRF center. To pool results from cells recorded with different saccade magnitudes, we normalized the final remapping magnitude by the saccade magnitude.

#### pRF size and eccentricity during perisaccadic remapping

Since the remapping direction becomes predominant forward starting about 25 ms before the saccade onset and the remapping process completes about 100 ms after the saccade offset (or about 150 ms after the saccade onset) ^6,10^, we focused on this period and divided it into four consecutive, non-overlapping 50-ms time bins centered at -25, 25, 75 and 125 ms relative to the saccade onset to calculate the pRF size and eccentricity of a cell in each bin. The 50-ms bin size balanced two competing factors: A smaller bin would introduce more noise into firing rate calculations whereas a larger bin would make it harder to study changes of RF size and eccentricity during remapping. It is also important not to include periods after the remapping ended as they would bias the results toward no change. For subsequent analyses, we retained only the data that met two criteria: (1) the neural response within a given time bin was significantly higher than the baseline; and (2) the remapping direction fell within 30° of the saccade (forward) direction. We then computed the pRF size and center as explained above, and the pRF eccentricity as the distance between the pRF center and the fixation. We next computed the changes of the pRF size and eccentricity in each time bin relative to the same cell’s cRF size and eccentricity (see below). Finally, we fit a mixed linear-effects model ^27^ to the change of the pRF size as a function of the change of the pRF eccentricity separately for the LIP and FEF cells, with both the slope and intercept as random variables.

Note that if we simply plotted the pRF size as a function of the pRF eccentricity, the well-known dependence of the RF size on the eccentricity among *different* cells would dominate the plot, and we could not examine the relationship between the RF size and eccentricity of the *same* cells during the remapping process. This is why we computed the *changes* of the pRF size and eccentricity in each time bin relative the same cell’s cRF size and eccentricity.

#### cRF size and eccentricity

As a control for the results in Fig. 7 (see text), we calculated the cRF size and eccentricity using responses from four 50-ms time bins centered at 75, 125, 175, and 225 ms after the probe onset, matching the time-bin size used in the pRF analysis. The first bin was centered at 75 ms after the probe onset because stimulus-evoked responses have a latency of about 50 ms. For each brain area, we randomly sampled a subset of the cRFs to match the corresponding pRF cell number in Fig. 7. We repeated this random subsampling 1000 times with replacement, and applied the robust linear regression ^28^ to the results of each subsample. We determined the mean and the 95% confidence interval of the slope from the distribution of fitted slopes. When we calculated the changes of a cell’s pRF size and eccentricity from the same cell’s cRF baselines, we used the cRF results from the 50-ms bin centered at 125 ms after the probe onset.

## Acknowledgement

This work was supported by US NIH (R01 EY032938) and National Natural Science Foundation of China (32030045 and 32061143004).

## Author contributions

NQ and YMW designed the study and the model. YMW simulated the model, reanalyzed the previously published data, and produced the figures. NQ, YMW, and MZ wrote the manuscript.

## Supplemental Information

### 1. Width of steady-state population activity bump

We first examined that when there is an activity bump evoked by a brief stimulus in our circuit model, what model parameters determine the width of the bump at the steady state. The analysis is similar to that of Amari et al who considered a step function as the activation function while we used a ReLU function (Equation 16). At the steady state after the brief input disappears, Equation 15 becomes:

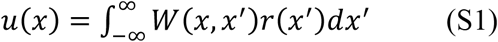

The population activity bump *r*(*x*) (or *u*(*x*) ≥ 0) is determined by the symmetric part of the connectivity, *W*_*sym*_(*x* − *x*^′^), and must have a symmetric shape. (The CD-gated anti-symmetric part, *W*_*anti*_, is only responsible for shifting the bump during saccades while maintaing its shape ^14^.) Without loss of generality, we can assume the bump extends from −*d* to +*d* with *u*(*x*) = *u*(−*x*). Since *u*(*x*) = *r*(*x*) for *u*(*x*) ≥ 0, Equation S1 becomes:

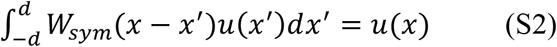

for −*d* ≤ *x* ≤ *d*, and *u*(*x*) = 0 otherwise. In particular, *u*(*x*) = 0 at *x* = −*d* and *d*:

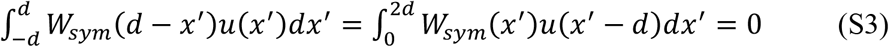

This equation determines the width of the activity bump 2*d*. As Fig. S1 illustrates, when *u*(*x*) is shifted by *d* to cover [0, 2*d*], its product with *W*_*sym*_(*x*) must have equal positive and negative areas to sum to 0. Since *W*_*sym*_(*x*) remains negative beyond the point of zero crossing, the width 2*d* is stable: under the perturbation *d* → *d* + *ε*, 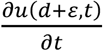 will have the opposite sign of *ε* to eliminate the perturbation. If the widths of *W*_*sym*_(*x*) and *u*(*x*) (and thus *d*) are scaled by the same factor, Equation S3 is still satisfied. This means that when *W*_*sym*_ is scaled to a different width, the activity bump (if present; see below) will scale accordingly.

For an activity bump *u*(*x*) to exist, Equation S2 implies that it must be an eigenvector of *W*_*sym*_ with an eigenvalue of 1. If we substitute: *W*_*sym*_(*x*) → *W*_*sym*_(*kx*), *u*(*x*) → *u*(*kx*) (and thus *d* → *kd*) to the left-hand side of Equation S2, we have:

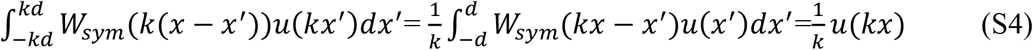

In other words, if *u*(*x*) is an eigenvector of *W*_*sym*_(*x*) with an eigenvalue of 1, then *u*(*kx*) is an eigenvector of *W*_*sym*_(*kx*) with an eigenvalue 1/*k*. Therefore, to have an eigenvalue of 1 for maintaining the activity bump, the proper substitution is *W*_*sym*_(*x*) → *kW*_*sym*_(*kx*). For example, if *k* = 2 to make *W*_*sym*_ (and the activity bump) half as wide, its strength should be doubled. Doubling the connection strength is equivalent to doubling the cell density.

The above results can be understood intuitively when the continuous equations are simulated with discrete neurons. Consider a circuit with N retinotopically arranged neurons that produces an activity bump of a certain stable size. We can assign the spacing between two adjacent neurons to any value of visual angle. If, for example, we half the assigned value, the width (in visual angle) of the activity bump will be halved and the neuron density (number of neurons per unit visual angle) will be doubled.

These considerations suggest that we can build the model in the cortical space and then assign a given cortical distance to a progressively larger visual angle as eccentricity increases according to Equation 1 in the text. To ensure the stability of the activity bump, *W*_*sym*_(*x, x*^′^) should be translationally invariant and symmetric in the cortical space *W*_*sym*_(*x, x*^′^) = *W*_*sym*_(*x* − *x*^′^) = *W*_*sym*_(*x*^′^ − *x*).

We finally note that since a stimulus evokes a symmetric population activity bump of size 2d in the cortical space, the RF of a cell has the same size of 2*d*, covering the range [*x-d, x+d*] for the cell tuned to *x*.

**Fig. S1.**
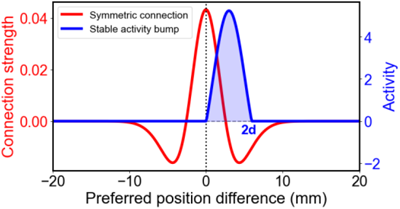
Stable population activity bump (blue) produced by the symmetric, Mexican-hat connectivity pattern (red). The width of the activity bump, 2*d*, in the cortical space is determined by the condition that the product of the two curves sums to 0.

### 2. Constant pRF sizes during remapping when the updating/remapping speed is uniform in the visual space

As we shown above, because the symmetric connectivity is uniform in the cortical space, the steady-state activity bumps and RFs are the same across the cortical space, covering a range of [-*d, d*] symmetrically around a given position *x*. Their shapes/sizes are different in the visual space and can be determined by Equation 1. For convenience, we define a cell’s RF center in the visual space as its peak location. Consider cell 1 in Fig. 4. In the cortical space, the left border, center, and right border of its RF are located at *x*_1_ − *d, x*_1_, and *x*_1_ + *d*, respectively (dashed arrows in Fig. 4a). In the visual space, they are located at *y*_1_ − *δ* = *E*(*x*_1_ − *d*), *y*_1_ = *E*(*x*_1_), and *y*_1_ + *μ* = *E*(*x*_1_ + *d*), respectively (dashed arrows in Fig. 4b), according to Equation 1, where:

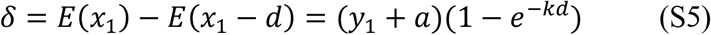

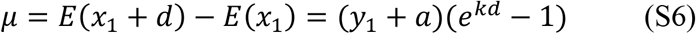

This means that small stimuli at *y*_1_ − *δ* and *y*_1_ + Δ produce population activity bumps that just include cell 1, and cell 1’s RF size in the visual space is Δ*y*_1_ = *δ* + *μ* (the distance between the dashed blue and green arrows in Fig. 4b).

Now consider cell 1’s RF during the CD-driven updating/remapping. Assume that at a given time *T* relative to the saccade onset, the activity bump evoked by a stimulus at *y*_2_ shifts to the bump centered at *y*_1_. Since the updating/remapping speed *v*_*y*_ is independent of *y*, at the same time *T*, the activity bump evoked by a stimulus at *y*_2_ − *δ* shifts to the bump centered at *y*_1_ − *δ*, and the activity bump evoked by a stimulus at *y*_2_ + *μ* shifts to the bump centered at *y*_1_ + *μ*. Since a steady-state activity bump is determined by the attractor dynamics and preserved by the anti-symmetric, CD-gated connections during the shift, it is the same at a given position *y* regardless of whether it is evoked by a stimulus at *y* or updated/remapped laterally to *y*. Therefore, when cell 1’s RF center is remapped from *y*_1_ to *y*_2_, its RF covers the range [*y*_2_ − *δ, y*_2_ + *μ*], with a size of Δ*y*_2_ = *δ* + Δ identical to its original size Δ*y*_1_.

### 3. Change of RF sizes during remapping when the updating/remapping speed is nonuniform in either the cortical space or the visual space

For the general case of *f*(*x*) = *f*(0)*e*^−*αkx*^ with 0 ≤ α ≤1, we have:

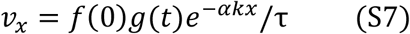

Then, Equations 1 gives:

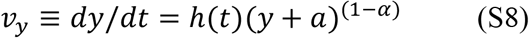

where *h*(*t*) = *kf*(0)*g*(*t*)/τ. Separate variables *y* and *t* and integrate over any fixed time window of duration *T* relative to the saccade onset during which an RF peaked at *y*_1_ is remapped forward to *y*_2_ (or equivalently, a population activity bump peaked at *y*_2_ is updated backward to *y*_1_), we have:

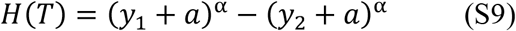

for α ≠ 0, and

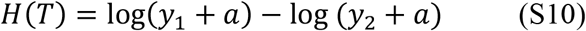

for α = 0. 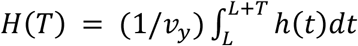 for an arbitrary but fixed starting time L and its detailed form is unimportant. In either case of α, we differentiate Equations S9 and S10 with respect to the *y* variables to obtain:

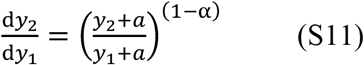

Equation S11 is exact. For finite sized Δ*y*_1_ around *y*_1_and Δ*y*_2_ around *y*_2_, the expression is approximate, which is Equation 14 in the text. The approximation happens to be exact for the two special cases of α = 0 and 1.

### 4. Data analyses with a different contour criterion

We used a standard contour criterion of 0.6 to produce Figs. 6 and 7 in the main text. To demonstrate the robustness of the results, we repeated the analyses with a contour criterion of 0.7. The results, shown in Figs. S2 and S3 are similar to those in Figs. 6 and 7.

**Fig. S2.**
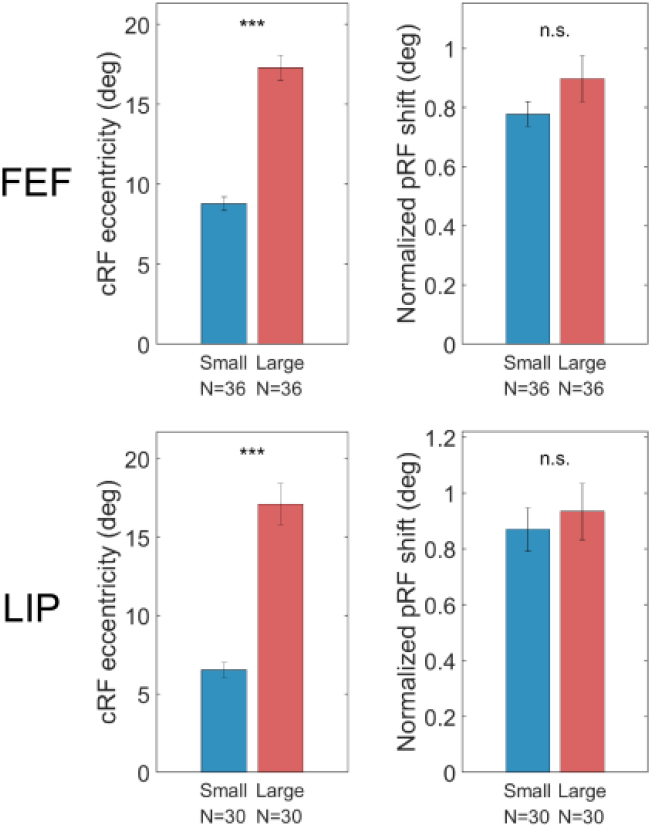
Spatially uniform remapping in LIP and FEF. The figure is identical to Fig. 6 of the main text except that we changed the contour criterion from 0.6 to 0.7. The left panels show that the small- and large-eccentricity groups of cells have significantly different mean eccentricities (two-sided Wilcoxon rank-sum test; p = 3.05×10^-13^ for FEF; p = 3.02×10^-11^ for LIP). The right panels show that the two groups have similar remapping magnitudes (p = 0.20 for FEF; p = 0.88 for LIP).

**Fig. S3.**
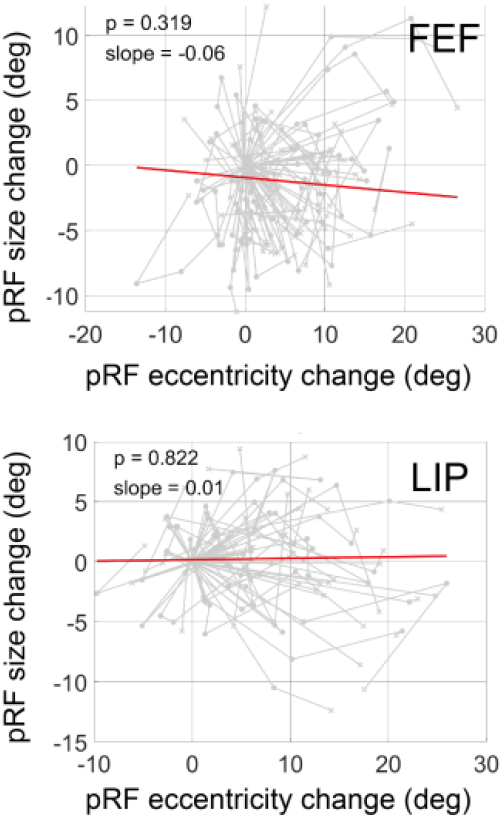
The change of pRF-size as a function of the change of eccentricity during remapping for cells in FEF (top) and LIP (bottom). The figure is identical to Fig. 7 of the main text except that we changed the contour criterion from 0.6 to 0.7. For each cell, the changes are relative to its cRF size and eccentricity during fixation (without remapping). The same cell’s results in different time bins (see text) are linked by gray lines. The cells numbers are the same as in Fig. S2. The red line in each plot is the fit of linear mixed model, which we use because different data points of the same cell are not independent samples.

